# Reproductive biology of the mudskippers *Boleophthalmus boddarti* and *Periophthalmus novemradiatus* in the Indian Sundarbans

**DOI:** 10.1101/2023.10.06.561162

**Authors:** Ishita Ray, Benjamin E. Himberg, Sanghita Sengupta, Sumit Homechaudhuri

**Affiliations:** Aquatic Bioresource Research Laboratory, Department of Zoology, University of Calcutta, 35 Ballygunge Circular Road, Kolkata, 700019, West Bengal, India; Department of Physics, University of Vermont, 82 University Place, Burlington, 05405, Vermont, United States of America (USA); Materials Science Program, University of Vermont, 82 University Place, Burlington, 05405, Vermont, United States of America (USA); Donostia International Physics Center (DIPC), Manuel Lardizabal Ibilbidea, 4, Donostia-San Sebastián, 20018, Spain

**Keywords:** Size at 50 % maturity, Fecundity, Spawning, Gonadosomatic index, Histology

## Abstract

Reproductive biology of mudskippers *Boleophthalmus boddarti* and *Periophthalmus novemradiatus* are investigated along the mudflats of the Indian Sundarbans. We investigated a total of 794 sampes for both male and female *Boleophthalmus boddarti* and *Periophthalmus novemradiatus* fish species to explore properties such as the size-at-50% maturity, fecundity, spawning, gonadosomatic and hepatosomatic indices along with the characterization of the macroscopic and histological attributes of the female gonads. Our work reports for the first time the weight-length relationships for these fish species in the Sundarbans region depicting a negative allometric growth pattern. The size-at-50% maturity which is often a crucial parameter for stock and reserve management is found for the female and male *Boleophthalmus boddarti* as 10.2 cm and 9.9 cm respectively, whereas, for the *Periophthalmus novemradiatus*, the same were 4.0 cm and 4.4 cm, respectively. Fully matured female *Boleophthalmus boddarti* were found to be multiple spawners in comparison to *Periophthalmus novemradiatus* with monthly variation of Le Cren’s relative condition ≤1, indicating good physiological conditions of the fish. The relationships between batch fecundity (F), body weight (i.e., BW, gm) and total length (i.e., TL, cm) were Log_10_F = 1.12 Log_10_BW + 2.41 and Log_10_F = 1.93 Log_10_TL + 1.82, respectively for *Boleophthalmus boddarti* and for the *Periophthalmus novemradiatus*, it was Log_10_F = 2.15 Log_10_BW + 2.57 and Log_10_F = 5.43 Log_10_TL - 0.81. The reproductive parameters estimated in this study are important information for studying population dynamics and stock assessments of these mudskippers in the Indian Sundarbans and will contribute to the management, conservation and prediction of environmental changes as these mudskippers are often studied as bio-indicators for climate-change effects.

## 1. Introduction

Fish reproductive biology plays an important role in aquaculture owing to its significance in stock assessment and fishery management. Crucial markers such as the weight-length relationships, condition factor, fecundity, somatic indices and the length at first maturity characterize the general reproductive patterns and well-being for a specific fish species. Often these markers are dependent on the particular environment and habitat of the fish and hence it is pertinent for us to understand and catalog such parameters for a given fish species within a certain habitat for optimum fishery management. This brings us to the topic of our work.

In this work, we investigate in detail the reproductive biology of the *Boleophthalmus boddarti* and *Periophthalmus novemradiatus* in the Indian Sundarbans. The Sundarbans Estuarine System (SES) is India’s largest monsoonal, macro-tidal, delta-front estuarine system located at the southernmost part of the Ganga-Brahmaputra delta bordering the Bay of Bengal. Additionally, the estuarine environment of the Sundarbans Biosphere Reserve is considered as a world heritage site by International Union for Conservation of Nature (IUCN) Mukher-jee et al. (2013) and has been inscribed as the “Wetlands of International Importance” Roy et al. (2020). It is very well-known that this mangrove estuarine ecosystem holds some of the most unique ecological properties Chatterjee et al. (2021); Ghosh et al. (2010) and serves as a novel habitation and foraging ground for numerous estuarine-based fish species. Although the Sunderbans serves as the nursery for nearly 90% of the aquatic species in the eastern coast of India, it faces enormous threat due to climate changes. Not to mention the severity of the reduction of freshwater discharge during lean seasons, increased salinity, use of destructive fishing gear, over-exploitation, extraction of resources and pollution Habib et al. (2020), all add to the risks associated with fishery management in this region.

Within the Sundarban estuarine system, the *Boleophthalmus boddarti* and *Periophthalmus novemradiatus* are abundant fish species which are widely distributed in the region’s inter tidal mudflats. They build burrows in the mudflats and during lowtides, when the burrows are exposed, the fish can be readily seen. In addition, they serve as distinctive ecological indicator in monitoring the estuarine and mudflat ecosysytem Ansari et al. (2014). Demographics of these fishes have been reported to comprise the areas spanning Diamond harbor (22.113°N and 88.115°E) and Namkhana (21.464°N and 88.132°E), West Bengal, India Mahadevan et al. (2021).

Let us mention that while an extensive study exists for the *Boleophthalmus boddarti* in the Mekong delta Nguyen et al. (2015) and Mumbai creeks Chandran et al. (2019), similar details about the reproductive patterns of this fish species is absent in the Indian Sundarbans and the general information about its reproductive biology is also rather scarce. Our work thus aims to bridge this gap and reports for the first time the parameters and markers that characterize the reproductive biology of *Boleophthalmus boddarti* in the Indian Sundarbans. In addition, to the best of our knowledge similar information on *Periophthalmus novemradiatus* has not been reported in the Indian Sundarbans yet. Thus this work is of significance in understanding not only the reproductive ecology but also provides with the necessary details that will be valuable for the conservation and management of these fish species.

The remainder of the paper is organized as follows: in Sec 2 we provide with the general methodology for the computation of the various parameters that characterize the reproductive biology and mention the details pertaining to the sample collection as well as the area of study. In Sec. 3 we lay out our results and finally end with a summary of these results and their significance in Sec. 4 .

## 2. Materials & Methods

### 2.1. Study site & Sample Collection

We conducted our study from March 2019 to February 2020 in the mudflats of Jharkhali island of Indian Sundarbans (Fig. 1). The island is located under the Basanti block of South 24 Parganas district in West Bengal. This island is considered a mid-estuarine region situated at the heart of the Sundarban biosphere reserve. It is surrounded by river Bidyadhari on the east, Matla on the west, and Herobhanga on the south. The region is characterized by a web of tidal water systems. The blue-spotted mudskipper and the Indian Dwarf mudskipper are resident species of this estuarine habitat. The fishes were collected from four sampling sites which were selected on the basis of habitat homogeneity. Being mudflats these sampling sites are completely inundated during high tide and are exposed during low tide. The mudskippers were collected during low tide when their burrows were visible in the mudflats. They were collected by various methods like handpicking and use of scoop net (0.5-2 cm mesh). The services of local fishermen were employed in the collection of the mudskippers. The fishes were identified using standard identification methods Ard et al. (1989); Polgar et al. (2013); Jaafar and Murdy (2017). The male and female of *Boleophthalmus boddarti* were distinguished on the basis of external morphology of their genitalia with oval relating to the female and triangular to the male Nguyen et al. (2015). The genital papilla of female *Periophthalmus novemradiatus* was broad and rounded whereas in males it is tapered and tongue-shaped Udo (2002). After collection, the fishes were measured for total length (TL nearest 0.1 cm) and body weight (BW nearest 0.01 gm). Gonads were collected from the fish samples and were packed with ice before laboratory processing.

**Figure 1:**
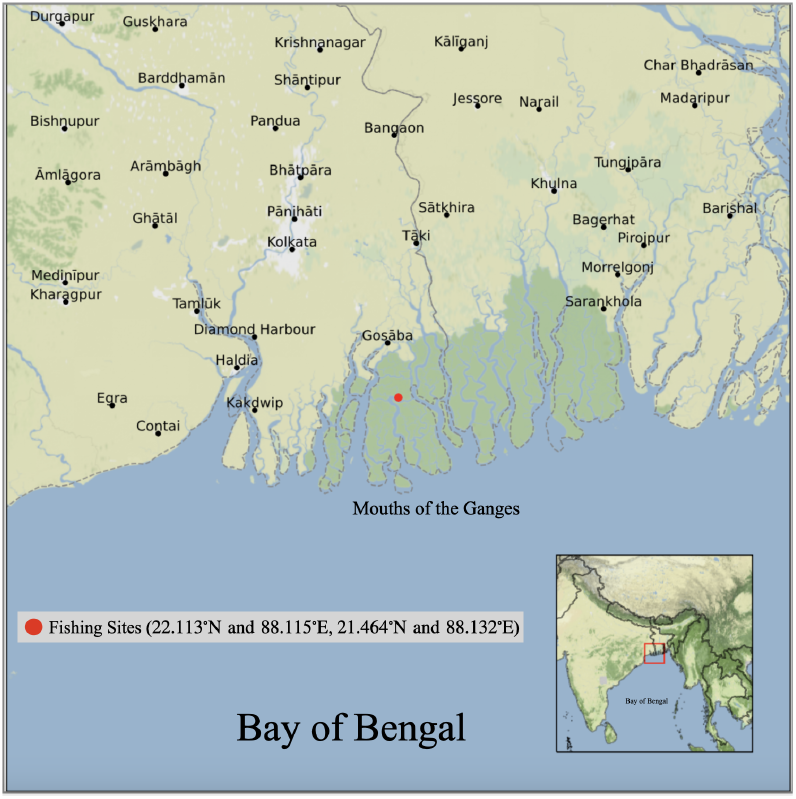
Location of the study area around the Indian Sundarbans with sample collection sites highlighted with red marker.

### 2.2. Laboratory Processing & Histological examination

Ovarian samples were weighed to the nearest 0.1 gm and preserved in 10 % buffered formalin solution for later histological examinaton. Fixed ovarian pieces were embedded in paraffin wax, sectioned at 4 micrometers, and stained/counterstained with hematoxylin/eosin following standard histological procedures. Based on the macroscopic appearance, ovaries are assigned with maturity stages as can be seen in Tables 1 and 2. The macroscopic and histological descriptions are provided in the slides for both the female *Boleophthalmus boddarti* and *Periophthalmus novemradiatus* along with corresponding details in respective Tables 1 and 2.

**Table 1:** Macroscopic stages and corresponding histological phases of ovarian development in female *Boleophthalmus boddarti*. Histological identification of different ovarian stages are shown in transverse sections with Stages I-V depicting development of ovary. *PO: Perinucleolus oocytes, CA: Cortical Alveolar oocytes, RO: Ripe oocytes, N: Nucleus, AF: Atretic follicle, EF: Empty follicle*.

| Stages | Macroscopic | Histological description |
| --- | --- | --- |
| I (Immature) | The ovaries are small, thin, elongated, yellowish white in color, vascularization is low. | Section shows presence of primary oocytes as perinucleolar oocytes. |
| II (Maturing) | Ovaries appear bigger, round and yellowish in color, blood vessels appear on the surface of the ovaries. | Section shows presence of cortical alveolar oocytes. Primary and secondary vitellogenic oocytes seen. |
| III (Mature) | Ovaries appear bigger, swollen, reddish yellow in color with prominent anterior, middle, posterior lobes. Eggs are translucent and can be seen on the surface of ovaries. | Section shows presence of larger cortical alveolar oocytes. Mature oocytes have higher yolk deposition. |
| IV (Ripe) | Ovaries are highly swollen, big in size, occupying almost entire abdominal cavity. Anterior, middle, posterior lobes are very much swollen, curved. Ovarian walls are thick and firm with prominent blood vessels. | Section shows presence of ripe oocytes with prominent nucleus and heavy yolk deposition. |
| V (Spent) | Ovaries are thin and contracted. Anterior, middle and posterior lobes have curly margins. Ovary appears yellowish orange in color with less vascularization. | Section shows presence of atretic follicles and empty follicles. Yolk granules have reduced in volume inside the oocytes. |

**Table 2:** Macroscopic stages and corresponding histological phases of ovarian development in *Periophthalmus novemradiatus*. Histologicalidentification of different ovarian stages are shown in transverse sections with Stages I-V depicting development of ovary. Stages I-V of development of ovary. *PO: Perinucleolus oocytes, CA: Cortical Alveolar oocytes, VO: Vitellogenic oocytes*.

| Stages | Macroscopic | Histological description |
| --- | --- | --- |
| I (Immature) | Ovary appears tiny, thread-like in appearance whitish in color | Transverse section shows the presence of pre-vitellogenic oocytes. Primary oocytes as perinucleolar oocytes are seen. |
| II (Maturing) | Ovary small, firm, pale and white. | Section shows presence of cortical alveolar oocytes. |
| III (Mature) | Ovary appears swollen, vascularized, orangish yellow, yellowish oocyte visible to naked eye through ovary wall. Ovary occupies 60 - 80 % of abdominal cavity. | Section shows presence of larger cortical alveolar oocytes. Cytoplasm shows small lipid vesicles. |
| IV (Ripe) | Ovaries appear very much swollen, turgid and highly vascularized, ovary wall transparent and eggs clearly distinct, ovary is deep yellow in color. | Section shows presence of larger sized granular oocytes consisting of yolk protein granules which are termed as vitellogenic oocytes. |
| V (Spent) | Ovaries are reduced in size, flaccid in appearance, no granulation, color is pale yellow, residual oocytes were present and visible through flaky wall. | Section shows presence of degenerating atretic oocytes; meshwork of connective tissue with few numbers of empty follicles. |

### 2.3. Weight-Length Relationships

The weight-length statistical data provides a cornerstone for fisheries management and often is used as a marker for rate of growth (change in weight), rate of reproduction as well as rate of mortality (Froese (2006); Bagenal (1957b); Jisr et al. (2018)). For the samples collected we measured the length (TL) and the body weight (BW) of the fish and mapped it to a curvilinear model given by BW = a ×TL^*b*^ to derive the constants *a* (*initial* growth co-efficient) and *b* (growth co-efficient) (Ricker (2010); Froese (2006)). Here, *b* is an exponent that varies among different fish species and describes the growth pattern in a fish. *Isometric* growth patterns (fish grows with an unchanging body form and unchanging specific gravity) are characterized by *b* = 3, whereas *allometric* growth pattern is shown by *b* ≠ 3 Froese (2006). Within the allometric pattern, there can be two cases where *b* < 3 which represents a fish with less girth as the length increases and *b* > 3 which shows increasing plumpness of the fish with increasing length Froese (2006).

### 2.4. Fish Body indices & Le Cren Condition Factor

We also investigate the Gonado-somatic index (GSI) and the Hepato-somatic index (HSI) which have been widely used to evaluate the timing of reproduction Lowerre-Barbieri et al. (2011). GSI serves as a metric that represents the ratio of fish gonad weight (GW) to the body weight (BW) given by, GSI = GW/BW × 100. Whereas, Hepatosomatic index (HSI) is the weight of the liver (LW) expressed as a percentage of total body weight given by, HSI = LW/BW ×100. The variation of HSI often reflects vitellogenesis in the liver Mousavi-Sabet et al. (2017). Additionally, using the co-efficients from the weight-length relationship, we will describe the *condition factor* of the fish. The index of well-being of the fish is often represented via Fulton’s condition factor, K = 100 × BW/TL^3^, where BW and TL (measured in cm) are the weight and length of the fish, respectively Fulton (1904); Mir et al. (2012); Sarkar et al. (2013). Fulton’s definition assumes that the weight and length of the fish increase isometrically (*b* = 3). On the other hand, Le Cren introduced the relative condition factor RCF = BW/(a ×TL^*b*^) Cren (1951) which does not assume the isometric growth pattern but is more suitable for fish species that exhibit allometric growth pattern (*b* ≠ 3). In this case, the observed body weight BW is divided by its predicted weight obtained from the linear regression of the weight-length relationship (BW = a ×TL^*b*^) Cren (1951).

### 2.5. Size at maturity

Size at maturity was treated statistically, with two stages of maturity classified as either immature PM=0 or mature PM=1 where PM is proportion mature (Ruiz Abierno et al. (2021); Okochi et al. (2016)). Maturity classification assigned by consideration of length, total weight and gonad state through dissection. For L_50_, the median length when half of the individuals reached maturity, a binary logistic regression was performed with the following equation:

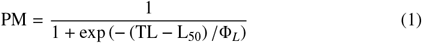

where TL is total length and Φ_*L*_ characterizes how sharply the curve transitions from immature to mature. The same binary logistic regression was performed for W_50_, the median body weight when half of the individuals reached maturity, by replacing TL with BW for body weight and Φ_*L*_ with Φ_*W*_ .

### 2.6. Batch fecundity

Ripe ovaries (Stage IV) were considered for the analysis of fecundity using the gravimetric method for fecundity estimation Bagenal (1957b,a). Sub samples measuring 0.1 gm were obtained from the anterior, middle and posterior parts of each formalin-fixed ovary. These sub samples were kept in Gilson fluid on the day of estimation to remove exterior connective tissues from the surface of the ovaries. Then the number of eggs were calculated from each sub sample and the mean number of eggs were calculated and then multiplied by total ovarian weight to get fecundity. The mean number of eggs between sub samples were converted into batch fecundity based on total gonad weight. When the coefficient of variation between the sub samples was higher than 10%, extraction of sub samples were conducted until the coefficient of variation was less than 10%.

### 2.7. Data Analysis

All statistical analyses were conducted in python-based code (PBCAF) Himberg et al. (2023).

## 3. Results

### 3.1. Weight-Length Relationships

We investigated the weight-length relationships for the female and male fish samples belonging to the *Boleophthalmus boddarti* and *Periophthalmus novemradiatus*, respectively (Fig. 2).

**Figure 2:**
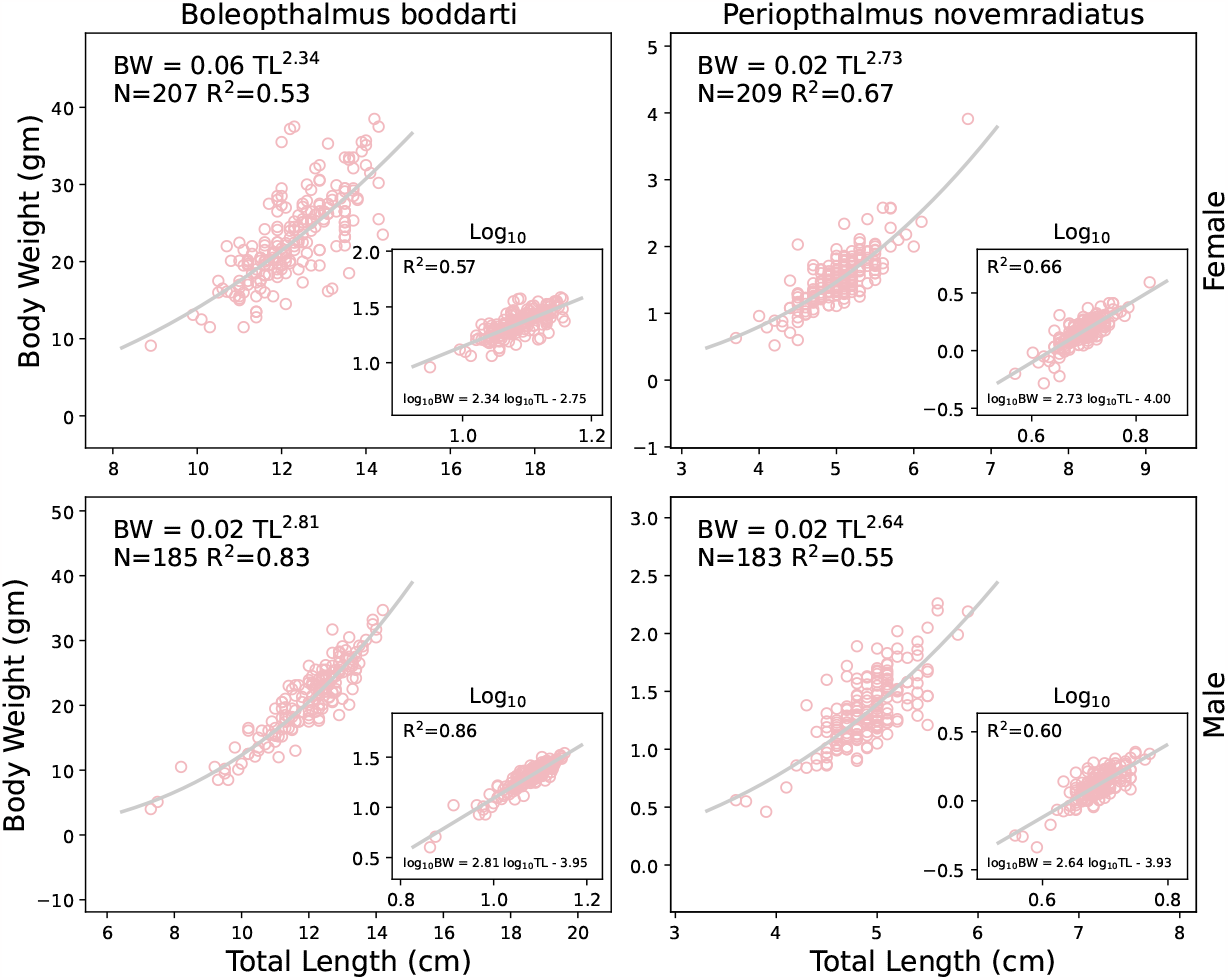
Weight-Length relationships for *Boleophthalmus boddarti* and *Periophthalmus novemradiatus. Upper panel: Female fish* and *Lower panel: Male fish. Insets* show the linear regressions of the Log(Body weight) *vs* Log (Total Length).

A total of 392 samples of *Boleophthalmus boddarti* were collected of which 207 were females and 185 were males. The range of length of female fishes was from 8.9 cm to 14.4 cm with an average of 12.3 ±0.98 cm. Whereas for the male fishes the range was 7.3 cm to 14.2 cm with an average 12.05 ±1.5 cm. We also measured their respective body weights. For the female *Boleophthalmus boddarti*, the body weights ranged from 9.1 gm to 38.5 gm with an average of 23.2 ±5.6 gm. For the male *Boleophthalmus boddarti*, the average bodyweight was 21.3 ±5.4 cm and the general range was 4.0 gm to 34.7 gm.

Similarly for the *Periophthalmus novemradiatus*, we collected 392 samples of which 209 were females and 183 were males. Compared to the *Boleophthalmus boddarti*, we observed that the *Periophthalmus novemradiatus* were much smaller in length as well as body weight. The average length and weight of the female *Periophthalmus novemradiatus* was measured as 5.01 ±0.38 cm and 1.51 ±0.38 gm, respectively. Their length ranged from 3.7 cm to 6.7 cm whereas the body weight was measured to range from 0.52 gm to 3.91 gm. For the males, we measured the average length and weight as 4.9 ±0.33 cm and 1.34 ±0.3 gm, respectively with the ranges for length being 3.6 cm to 5.9 cm and body weight as 0.46 gm to 2.26 gm.

In case of the female *Boleophthalmus boddarti*, we found BW = 0.06 TL^2.34^ (N = 207, R^2^ = 0.53), whereas for the female *Periophthalmus novemradiatus*, the relationship was calculated as BW = 0.02 TL^2.73^ (N = 209, R^2^ = 0.67). For the males, we estimated BW = 0.02 TL^2.81^ (N = 185, R^2^ = 0.83) for the case of *Boleophthalmus boddarti* and BW = 0.02 TL^2.64^ (N = 183, R^2^ = 0.55) for *Periophthalmus novemradiatus*, respectively. In the insets of Fig. 2, we additionally show the variation of the logarithm of the length of the fish (Log_10_ TL) with the logarithm of the body weight of the fish (Log_10_ BW) with the respective regression expressions. From the weight-length relationships, we note that both the fish species show negative allometric growth pattern (*b* < 3) and thus is rather slender.

### 3.2. Size at maturity

Size at maturity was analyzed for female and male fish samples belonging to the *Boleophthalmus boddarti* and *Periophthalmus novemradiatus* both as a function of total length TL and body weight BW respectively (Fig. 3 and Fig. 4).

**Figure 3:**
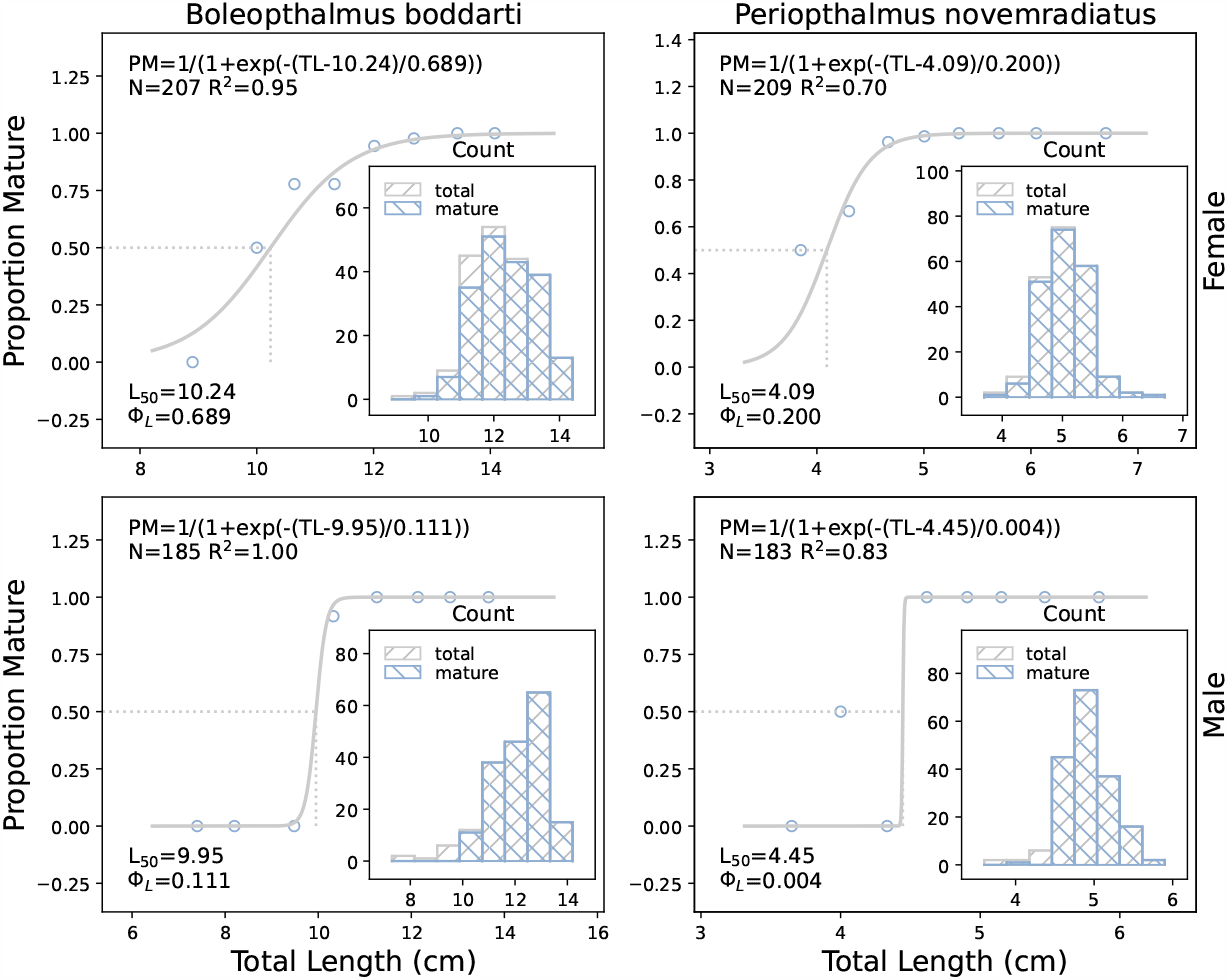
Sigmoid curve showing length at first maturity of reproductive female (*upper panel*) and male (*lower panel*) for *Boleophthalmus boddarti* and *Periophthalmus novemradiatus. Insets* show histogram of length-frequency distribution in collected samples as also shown in Table 4 and Table 3.

**Figure 4:**
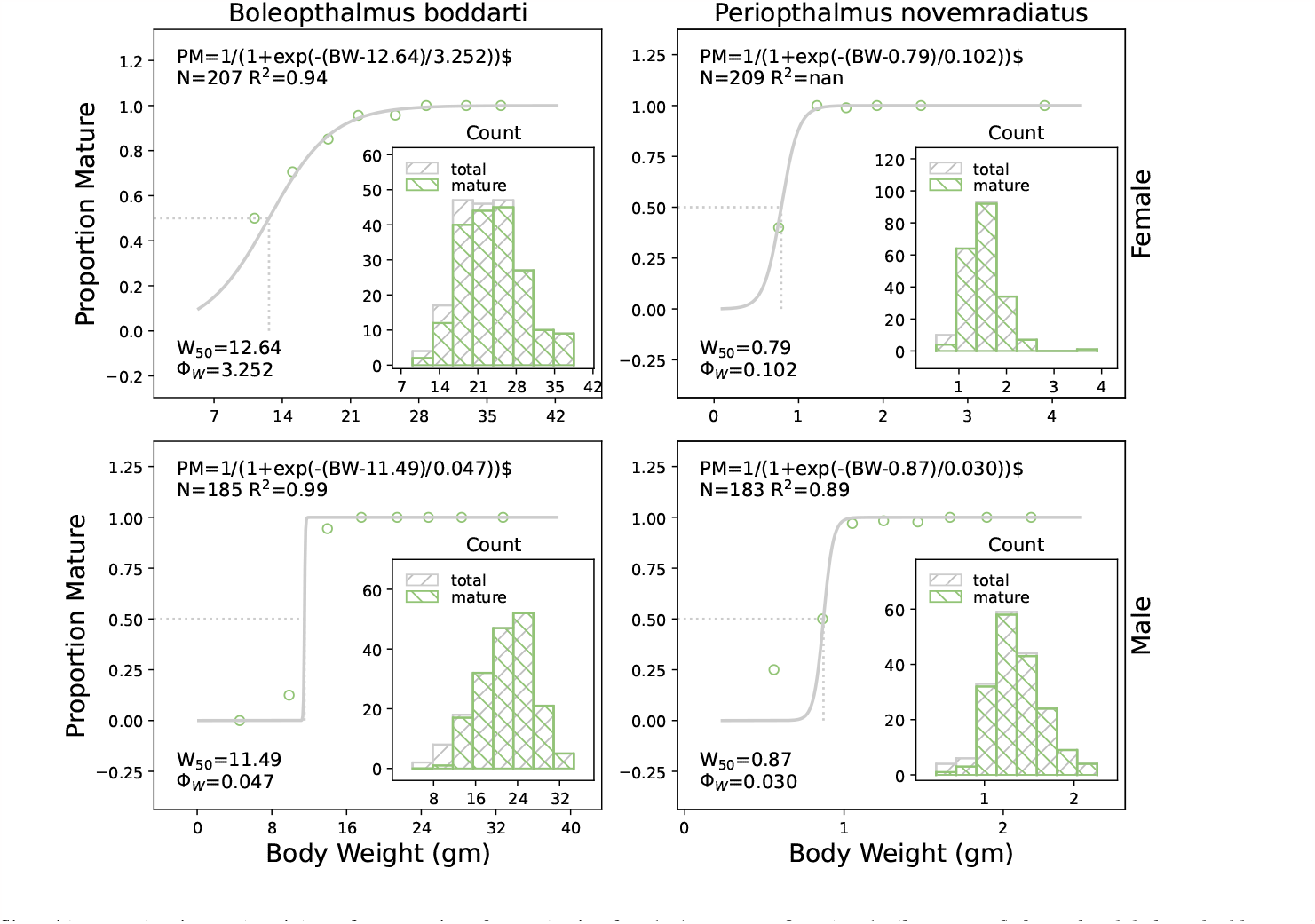
Sigmoid curve showing bodyweight at first maturity of reproductive female (*upper panel*) and male (*lower panel*) for *Boleophthalmus boddarti* and *Periophthalmus novemradiatus. Insets* show histogram of body weight-frequency distribution in collected samples as also shown in Table 6 and Table 5 respectively.

A total of 392 specimens were dissected for *Boleophthalmus boddarti*, consisting of 207 females and 185 males. The same number of specimens were processed for *Periophthalmus novemradiatus*, however there were 209 females and 183 males. A discussion of the observed BW and TL for both *Boleophthalmus boddarti* and *Periophthalmus novemradiatus* can be found in Sec. 3.1. Each specimen was additionally assigned a maturity of 0 for immature or 1 for mature. This maturity was determined by inspection of TL, BW and also dissection to inspect the state of the gonads.

**Table 3:** *Boleophthalmus boddarti* average total length (TL) and maturity data used to determine L_50_, as shown in the *left panel* of Fig. 3. The first column of the table is the average total length for each bin in the histogram, and the second column is the standard deviation (SD) of samples used to determine the average.

| Female |  |  |  |
| --- | --- | --- | --- |
| TL Average (cm) | TL SD (cm) | Mature | Total |
| 8.90 | - | 0 | 1 |
| 10.00 | 0.10 | 1 | 2 |
| 10.64 | 0.18 | 7 | 9 |
| 11.33 | 0.22 | 35 | 45 |
| 12.01 | 0.17 | 51 | 54 |
| 12.69 | 0.19 | 43 | 44 |
| 13.44 | 0.19 | 39 | 39 |
| 14.08 | 0.20 | 13 | 13 |

Table 3: *Boleophthalmus boddarti* average total length (TL) and maturity data used to determine $L_{50}$ , as shown in the *left panel* of Fig.
| Male |  |  |  |
| --- | --- | --- | --- |
| TL Average (cm) | TL SD (cm) | Mature | Total |
| 7.40 | 0.10 | 0 | 2 |
| 8.20 | - | 0 | 1 |
| 9.48 | 0.20 | 0 | 6 |
| 10.33 | 0.23 | 11 | 12 |
| 11.26 | 0.24 | 38 | 38 |
| 12.14 | 0.21 | 46 | 46 |
| 12.84 | 0.26 | 65 | 65 |
| 13.66 | 0.26 | 15 | 15 |

**Table 4:** *Periophthalmus novemradiatus* average total length (TL) and maturity data used to determine L_50_, as shown in the *right panel* of Fig. 3. The first column of the table is the average total length for each bin in the histogram, and the second column is the standard deviation (SD) of samples used to determine the average.

| Female |  |  |  |
| --- | --- | --- | --- |
| TL Average (cm) | TL SD (cm) | Mature | Total |
| 3.85 | 0.15 | 1 | 2 |
| 4.30 | 0.11 | 6 | 9 |
| 4.67 | 0.11 | 51 | 53 |
| 5.00 | 0.08 | 74 | 75 |
| 5.33 | 0.12 | 58 | 58 |
| 5.70 | 0.09 | 9 | 9 |
| 6.05 | 0.05 | 2 | 2 |
| 6.70 | - | 1 | 1 |

Table 4: *Periophthalmus novemradiatus* average total length (TL) and maturity data used to determine $L_{50}$ , as shown in the *right panel* of Fig.
| Male |  |  |  |
| --- | --- | --- | --- |
| TL Average (cm) | TL SD (cm) | Mature | Total |
| 3.65 | 0.05 | 0 | 2 |
| 4.00 | 0.10 | 1 | 2 |
| 4.33 | 0.07 | 0 | 6 |
| 4.62 | 0.08 | 45 | 45 |
| 4.91 | 0.08 | 73 | 73 |
| 5.15 | 0.07 | 37 | 37 |
| 5.46 | 0.07 | 16 | 16 |
| 5.85 | 0.05 | 2 | 2 |

**Table 5:**
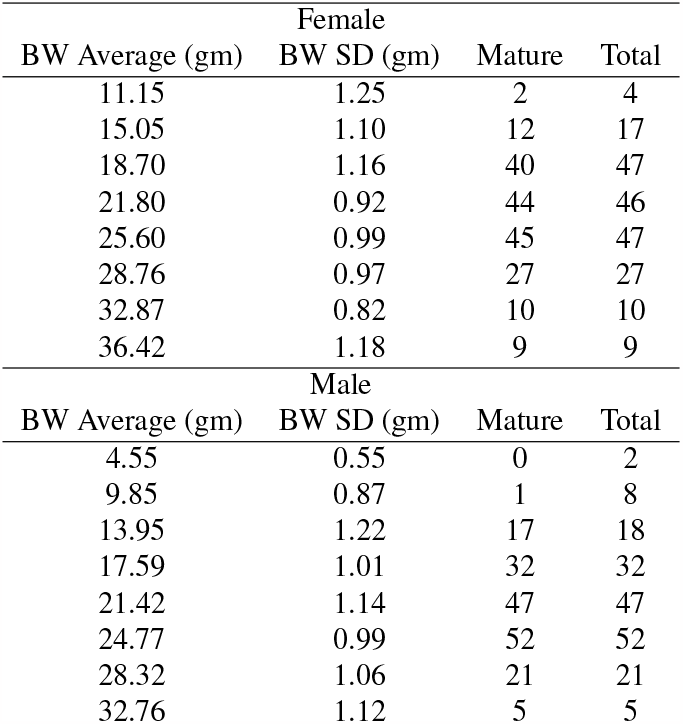
*Boleophthalmus boddarti* average body weight (BW) and maturity data used to determine BW_50_, as shown in the *left panel* of Fig. 4. The first column of the table is the average body weight for each bin in the histogram, and the second column is the standard deviation (SD) of samples used to determine the average.

**Table 6:** *Periophthalmus novemradiatus* average body weight (BW) and maturity data used to determine BW_50_, as shown in the *right panel* of Fig. 4. The first column of the table is the average body weight for each bin in the histogram, and the second column is the standard deviation (SD) of samples used to determine the average.

| Female |  |  |  |
| --- | --- | --- | --- |
| BW Average (gm) | BW SD (gm) | Mature | Total |
| 0.76 | 0.13 | 4 | 10 |
| 1.22 | 0.10 | 64 | 64 |
| 1.56 | 0.12 | 92 | 93 |
| 1.93 | 0.10 | 34 | 34 |
| 2.45 | 0.12 | 7 | 7 |
| - | - | 0 | 0 |
| - | - | 0 | 0 |
| 3.91 | - | 1 | 1 |
| Male |  |  |  |
| BW Average (gm) | BW SD (gm) | Mature | Total |
| 0.56 | 0.07 | 1 | 4 |
| 0.87 | 0.02 | 3 | 6 |
| 1.05 | 0.06 | 32 | 33 |
| 1.25 | 0.06 | 58 | 59 |
| 1.46 | 0.07 | 43 | 44 |
| 1.67 | 0.06 | 24 | 24 |
| 1.90 | 0.07 | 9 | 9 |
| 2.16 | 0.08 | 4 | 4 |

There were 175 *Boleophthalmus boddarti* males found to be mature, and 10 males found to be immature. There were 189 females found to be mature, and 18 females found to be immature. A binary logistic regression was performed for both PM vs TL and PM vs BW, as described in Sec. 2.5, where PM is proportion mature. For *Boleophthalmus boddarti*, it was found that L_50_ = 9.95 cm and L_50_ = 10.24 cm for the males and females, respectively. Similarly, it was found that W_50_ = 11.49 gm and W_50_ = 12.64 gm for the males and females respectively.

For both L_50_ and W_50_ binary logistic regressions, Φ_*L*_ and Φ_*W*_ were considerably smaller for males at Φ_*L*_ = 0.111 cm and Φ_*W*_ = 0.047 gm compared to the females Φ_*L*_ = 0.689 cm and Φ_*W*_ = 3.25 gm. This suggests a much steeper maturity curve for the males which in turn suggests that males, once they reach a specific body weight of total length, mature more quickly than females.

In the case of *Periophthalmus novemradiatus*, there were 174 males found to be mature and 11 males found to be immature. There were 202 females found to be mature and 7 females found to be immature. A similar binary logistic regression was performed, as with *Boleophthalmus boddarti*. For *Periophthalmus novemradiatus*, it was found that L_50_ = 4.45 cm and L_50_ = 4.09 cm for the males and females respectively. It was also found that W_50_ = 0.87 gm and W_50_ = 0.79 gm for the males and females respectively. For both L_50_ and W_50_ binary logistic regressions, Φ_*L*_ and Φ_*W*_ were smaller for males at Φ_*L*_ = 0.004 cm and Φ_*W*_ = 0.030 gm compared to the females Φ_*L*_ = 0.200 cm and Φ_*W*_ = 0.102 gm.

For both Fig. 3 and Fig. 4, R^2^ values were determined using the fitted curve and scatter points present in the plots. The scatter points were calculated from the histogram as the ratio of mature to total fish in each bin vs the average TL or BW for each bin. The averages and standard deviations (SD) of TL can be found in Table 4 for *Periophthalmus novemradiatus* and Table 3 for *Boleophthalmus boddarti*. Table 6 and Table 5 provide averages and standard deviations (SD) of BW for *Periophthalmus novemradiatus* and *Boleophthalmus boddarti* respectively.

### 3.3. Spawning Season & Monthly variation in mean fish Body indices

First, we begin with the description of the gonadosomatic index (GSI). For female *Boleophthalmus boddarti*, in Fig. 5 (upper panel left column), we find that GSI peaks in the early premonsoon for the months of February and March. The rise in GSI is continued till the end of pre-monsoon in May. From the beginning of monsoon *i.e* from June onwards, the GSI value is found to decrease indicating the beginning of spawning in the fishes. By the end of monsoon in September, the GSI value is seen to drop, pointing to the conclusion of spawning event in the fishes. From the beginning of post-monsoon, we find that the GSI again starts to increase and by the mid of post-monsoon, GSI drops indicating spawning event. Again the GSI value starts to increase from the premonsoon season repeating the cyclical development of the gonads. Thus the GSI values indicate that the fish is a multiple spawner and reproduces throughout the year. As a comparison let us point out that the study of reproductive parameters of *Boleophthalmus boddarti* in Ref. Nguyen et al. (2015) from March 2013 to February 2014 showed that ovary developed from immature to mature stage in between four months spanning from July to October and spent ovaries were found in the month of September. The temporal occurrence of the spent ovary from Ref. Nguyen et al. (2015) is seen to coincide with our findings. For the male *Boleophthalmus boddarti* we see a similar pattern of gonad development like the females. The GSI value peaks up in the start of the early monsoon and then begins to drop. This drop indicates the event of spawning. The GSI does not show much variation through out the monsoon season indicating that the gonad remains in spawning stage throughout the monsoon. Post spawning the GSI begins to rise, i.e the gonads start to grow. The GSI attains another peak in the begining of the post-monsoon and drops after that. Thus time peaks of GSI in this cycle indicates that the fish is a multiple spawner.

**Figure 5:**
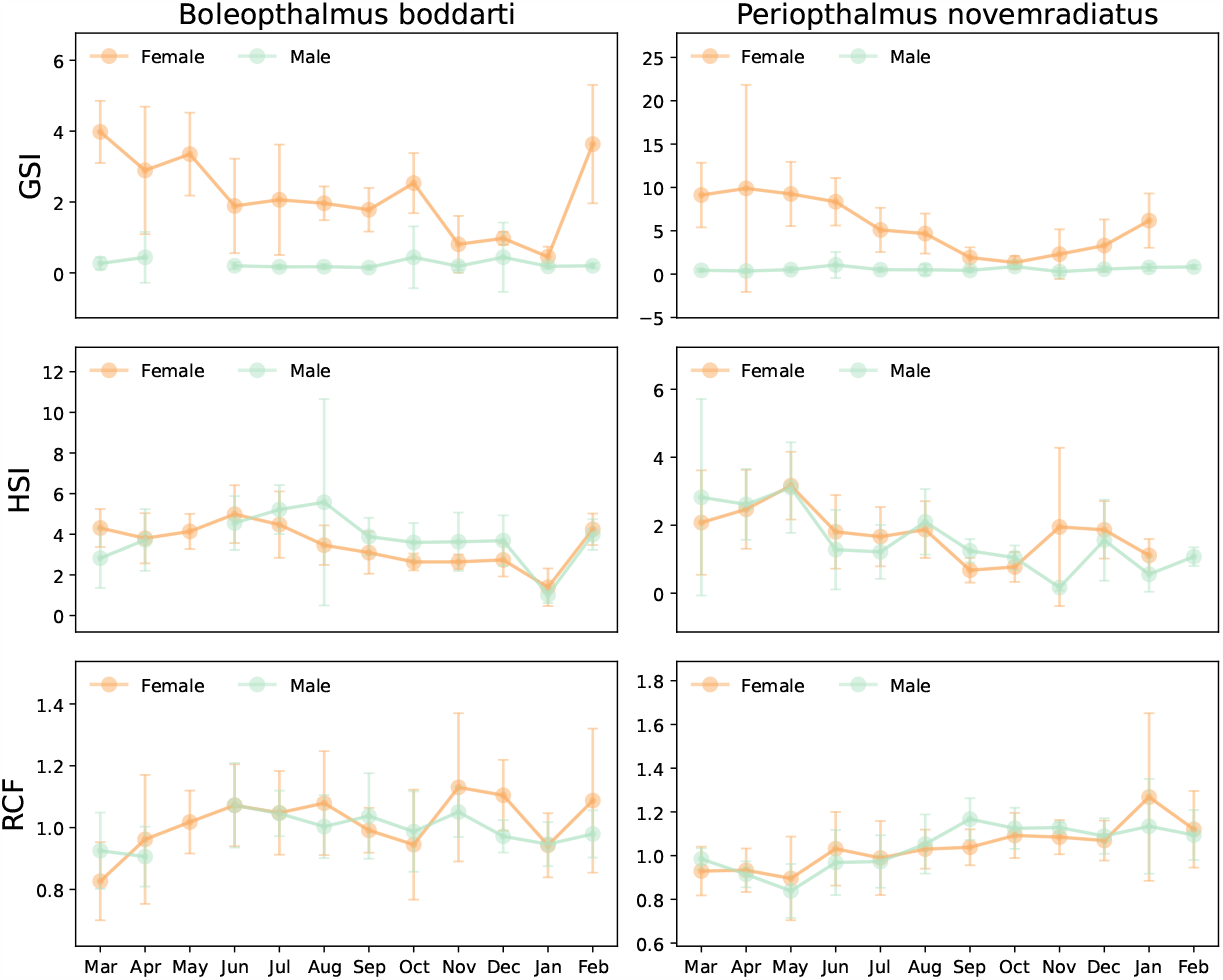
Comparative description of *somatic indices* and relative condition factor for *Boleophthalmus boddarti* and *Periophthalmus novemradiatus. Upper panel*: Mean gonadosomatic index (GSI) ± standard deviation with monthly variation for both male and female fish species.*Middle panel*: Mean hepatosomatic index (HSI) ± standard deviation with monthly variation for both male and female fish species. *Lower panel: Le Cren* condition factor (relative condition factor RCF) ±standard deviation with monthly variation for both male and female fish species.

Next, we provide some results on the hepatosomatic index (HSI) of the female *Boleophthalmus boddarti* also shown in Fig. 5. The hepatosomatic index value is seen to increase during the early ripening stages of the ovary in the beginning of premonsoon and then decreases in the late ripening stages when the fish starts to spawn in the monsoon. Thus the HSI value shows a similar trend with that of GSI with reaching peak in early pre-monsoon and post-monsoon when the ovaries begin to mature. During monsoon when the ovaries are in spent stage both the HSI and GSI drops. The HSI value of the mudskipper *Periopthalmus barbarus* studied in mangrove swamps of Iko estuary, South east Nigeria between November, 2011 to October, 2013 also showed this trend with the GSI value Udo (2002). The HSI value of *Periophthalmus barbarus* sampled over a twelve month period (February 2008 – January 2009) in New Calabar River Nigeria showed a gradual decrease from April to July with the GSI value also dropping during the same period Chukwu et al. (2010) indicating breeding activities during that period. For the male *Boleophthalmus boddarti*, the HSI value peaks up in the pre-monsoon end, supporting the gonad development phase during this time. At the begining of monsoon, the HSI begins to drop. During this stage, the gonads are also in spent stage. Again in the post-monsoon, a rise in HSI is seen. During this time, the liver supports the growth of the gonad past the spawning event. In the late post-monsoon, the HSI again drops and is then seen to rise.

In Fig. 5, right column we show the GSI and HSI of the female and male *Periophthalmus novemradiatus* as well. We first provide a discussion of the results for the GSI of the female fish. We find the GSI values to peak up from early pre-monsoon for the months March indicating the development of ovary for the event of spawning. From the begining of monsoon, i.e, June onwards, the GSI starts to decrease pointing to the event of spawning. The spawning event continues till the end of monsoon i.e, till September. Again the GSI value starts to increase from post-monsoon begining from October and continues till February which marks the begining of pre-monsoon. So the drop in GSI indicating the spawning event can be seen only in the monsoon season for this fish. Thus it is not a multiple spawner like the blue-spotted mudskipper. Spawning season for this fish spans from June to September. The HSI value of the female *Periophthalmus novemradiatus* is seen to increase during the early ripening stages of the ovary in the beginnign of the pre-monsoon and then decreases in the late ripening stages when the fish starts to spawn in the monsoon. During monsoon when the ovaries are in spent stage both HSI and GSI drops.

For the male *Periophthalmus novemradiatus* the gonad shows a steady state of growth throughout the breeding cycle/annual cycle. The GSI value of the male fish is also reflective of the above said fact. The GSI increases during the premonsoon with a peak just at the begining of the monsoon (June) and then it starts to drop off indicating begining of the spawning event. The spawning event spans throughout the monsoon season as seen by the drop of the GSI. The HSI of the male *Periophthalmus novemradiatus*, shows a rise when the gonads are developing during the pre-monsoon season and it starts to drop during the begining of monsoon. This drop in HSI occurs when the fish is in spawning stage of its breeding cycle during monsoon season. Again when the gonads are developing post the spawning event, the HSI begins to rise in the post-monsoon season.

In this section, we also show the results for the monthly variation of the *Le Cren condition factor* (RCF) of the fish using the co-efficients from the weight-length relationship. As the value of the co-efficient *b* < 3, thus relative condition factor is more appropriate for these fish species in comparison to Fulton’s condition factor. In the lower panel of Fig. 5, we show the RCF variation for these fishes. We note that the value of RCF ≤1 denote the good condition of the fish. High and low peaks characterize how the energy of the fish is invested in the growth of the gonad Hashim et al. (2017). In case of female *Boleophthalmus boddarti*, we observe 4 different peaks that correspond to months June, August, November and February. High condition factor corresponding to certain months is indicative of the energy of the fish being invested on growth after recovering from a spawning season.

### 3.4. Batch Fecundity

Fig. 6 shows relationship between fecundity and the total length and body weight of the fish (both characterized in logarithmic scales). We observe linear relationships between fecundity and length as well as fecundity and body weight of the fish. This suggests that with the increase in body weight and gonad weight of the fish, the number of eggs in the ovaries increases proportionally. This linear relationship is well-known and has been demonstrated by various studies Chandran et al. (2019); Nguyen et al. (2015). In Ref. Nguyen et al. (2015), the authors reported the batch fecundity of *Boleophthalmus boddarti* to be 9800-33800 eggs corresponding to a data collected during the time period March 2013 to February 2014. In our work, we find the batch fecundity of *Boleophthalmus boddarti* to vary between 4897-29024 eggs which is lower than that reported in the study of Ref. Nguyen et al. (2015). This difference could be related to the changes in the environmental factors affecting the habitat of the fish in the Mekong delta versus Sundarban regions.

**Figure 6:**
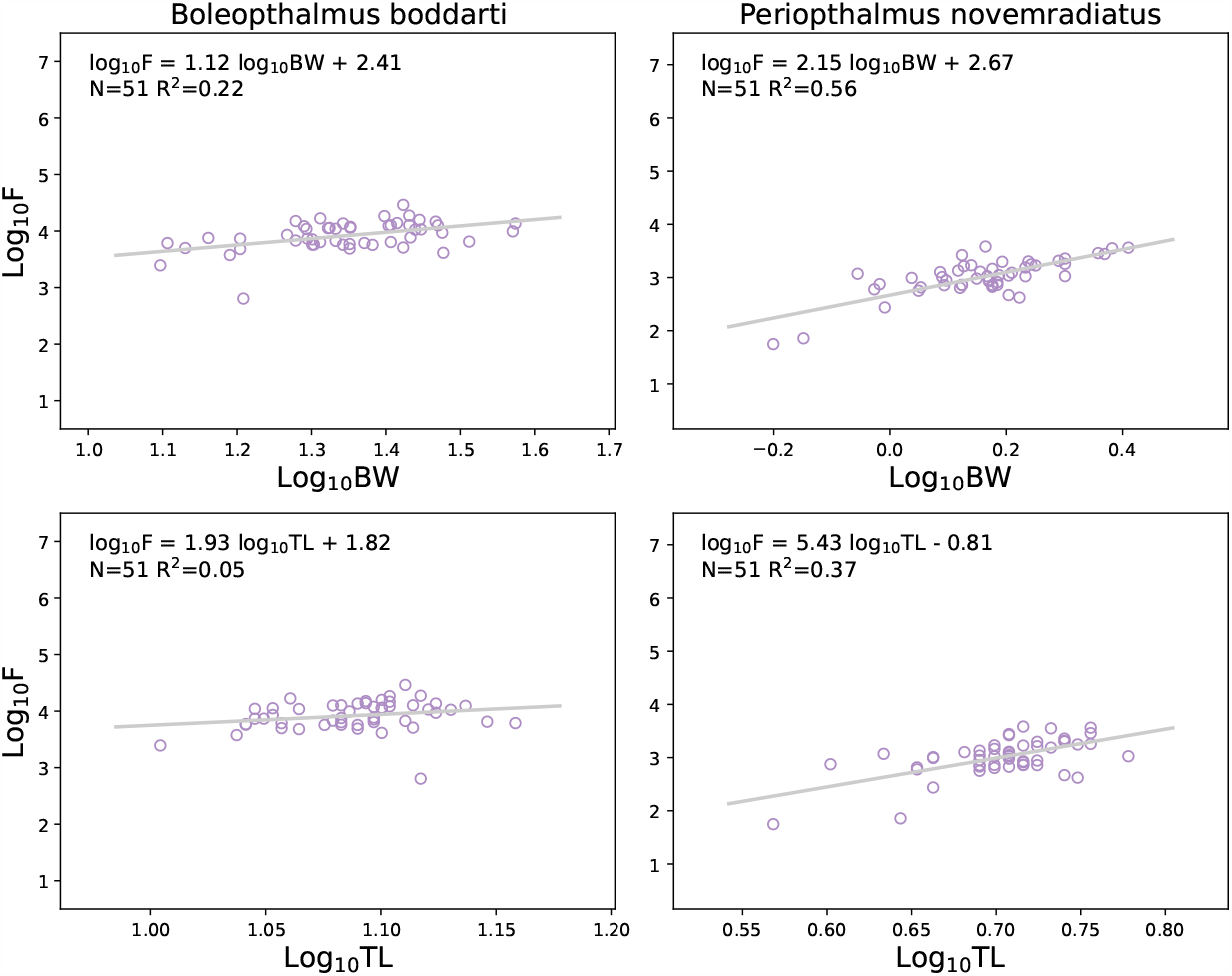
Fecundity variations in *Boleophthalmus boddarti* and *Periophthalmus novemradiatus. Upper panel:* Relationship between fecundity (Log F) and body weight (Log BW) of fish. *Lower panel:* Relationship between fecundity (Log F) and total length (Log TL) of fish.

The linear regressions as shown in the figures depict a positive relationship between the fecundity (Log F) and the total length of the fish (Log TL). Similar features are also observed for the relationship between fecundity and the body weight of the fish (Log TL vs Log BW). These computations were performed for *N* = 51 total number of fishes and the linear fits are characterized by R^2^ = 0.22 and R^2^ = 0.05 for the respective plots with expressions for regressions as Log_10_F = 1.12 Log_10_BW + 2.41 and Log_10_F = 1.93 Log_10_TL + 1.82. General trends such as these are also reported in Ref. Nguyen et al. (2015).

We investigated fecundity relationships also for the *Periophthalmus novemradiatus* and found positive relationship between the fecundity (Log F) and total length (Log TL) as well as body weight (Log BW). For a sample size of N = 51, we found Log_10_F = 2.15 Log_10_BW + 2.57 with R^2^ = 0.56. Meanwhile corresponding to the length, we found Log_10_F = 5.43 Log_10_TL - 0.81 with R^2^ = 0.37.

## 4. Conclusion

*Boleophthalmus boddarti* and *Periophthalmus novemradiatus* are both abundant fish species in the mudflats of the Indian Sundarbans. Information about their reproductive biology is scarce in the Indian subcontinent and especially in the adverse climatic conditions in the Sundarbans. In this work we have discussed many features related to the reproductive biology of both the fish species. Our results show that the fish species exhibits negative allometric growth pattern with *b* < 3 indicating the fish is more slender. Our work reports for the first time this metric for these mudskippers. We also provide with the information about the variation of the Le Cren’s relative condition factor (RCF) for these fishes and show its seasonal variation. Our results indicate that the physiological condition of both the fishes is good with RCF ≤1. We further provide with detailed relationships between the fecundity and the length as well as body weight of both the mudskippers. Our works are in match with general trends of fecundity as seen in *Boleophthalmus boddarti* species in the Mekong delta Nguyen et al. (2015). Slight variation in the parameters between the studies performed in the Mekong delta Nguyen et al. (2015) and our work in Sundarbans is indicative of the role of the habitat of the fish species. We also have performed histological studies that provide with the development of the ovary of these fishes and investigate in details the seasonal variation of the hepatosomatic and gonadosomatic indices which are crucial markers for fish reproductive biology. Lastly, we provide with the length (L_50_) and body weight (W_50_) at first maturity for these fishes and show detailed analysis for the proportion of maturity for both the male and female species. This value is of importance in fisheries management and stock assessment.

## Data availability

All accompanying data files used for the analysis will be made available upon request. Code for the data analysis will be made available to use at Github Himberg et al. (2023).

## Acknowledgements

SS was supported by the European Union (EU) through Horizon 2020 (FET-Open project SPRING Grant no. 863098). IR is grateful to the insightful discussions with the members of the Aquatic Bioresource Research Laboratory.

## Notes

### Competing Interest Statement

The authors have declared no competing interest.

